# SPEND-hSRS imaging of fumarate uncovers mitochondrial metabolic heterogeneity

**DOI:** 10.64898/2026.04.03.716311

**Authors:** Dingcheng Sun, Guangrui Ding, Haonan Lin, Guo Chen, Chun-Chin Wang, Seema Bachoo, Sarah E. Bohndiek, Ji-Xin Cheng

## Abstract

Mitochondria, acting as the energy powerhouse, biosynthetic center, and reductive equivalent hub of the cell, participate in cellular metabolic activities. However directly imaging mitochondrial chemical content and quantifying metabolic activity in living cells remain challenging. Here, by Self-PErmutation Noise2noise Denoiser enhanced Hyperspectral Stimulated Raman Scattering (SPEND-hSRS) microscopy, we demonstrate fingerprint-region metabolic imaging of fumarate, a key intermediate in the tricarboxylic acid (TCA) cycle, with sub-millimolar sensitivity. In chemotherapy-stressed bladder cancer cells, fumarate imaging revealed two mitochondrial subpopulations with divergent TCA metabolic preferences quantified by ratio metric analysis. Pixel-wise least absolute shrinkage and selection operator (LASSO) spectral unmixing further reconstructs fumarate and lipid maps, uncovering localized fumarate enrichment in protrusions. Extending to CH-window hyperspectral SRS imaging, we uncover the interplay between mitochondria and lipid droplets (LDs) in protrusions, where fatty acid is found to be released from LDs, to fuel the TCA cycle. Together, our work establishes SPEND-hSRS as high-resolution platform for linking fumarate to mitochondrial heterogeneity. Our results provide new insights into how mitochondrial heterogeneity and interaction with LDs drive cancer cell adaptation to stress.

## Main

Mitochondria serve as the energy metabolism hub of the cell, supporting ATP production through the tricarboxylic acid (TCA) cycle and oxidative phosphorylation ^1,2^. The metabolic demands of a cell vary by cell type and even among subcellular compartments^3,4^. Recent research reveals that mitochondria separate into two functional subpopulations based on the bioenergetic demand^5^. Thus, visualizing mitochondrial metabolism heterogeneity in live cells is key to understanding the machinery of life and diseases such as cancer^6^ and neurodegeneration^7^.

Current approaches to assess mitochondrial activity rely on metabolite- and protein-based strategies. Metabolite-based measurements most commonly rely on liquid chromatography–mass spectrometry (LC–MS). However, such approaches often combine with intracellular isotope labelling, requiring extensive sample processing, large cell numbers, and disrupt subcellular organization, preventing single-cell or organelle-level analysis^8,9^. Protein-based assays^10^, including fluorescent probes and genetically encoded biosensors^11–13^, provide indirect functional readouts but can perturb cellular metabolism and suffer from limited specificity or quantitative accuracy^14^.

To overcome these limitations, we developed SPEND-hSRS microscopy, which integrates a self-permutation Noise2Noise denoiser (SPEND)^15^ with hyperspectral SRS (hSRS), to map intracellular fumarate, a key TCA cycle intermediate (**Extended Data Fig. 1a**) through its intrinsic Raman signature at 1401 cm^-1^ (**Extended Data Fig. 1b**). While spontaneous Raman scattering microscopy is limited to ∼8 mM sensitivity^16^, our method pushes the imaging limit of detection (LOD) of fumarate into the sub-millimolar range, enabling high-fidelity downstream analysis, including pixel-wise LASSO spectral unmixing^17^.

By pharmacological TCA cycle blockade and SPEND-hSRS imaging, we directly visualized intracellular fumarate distributions across multiple cancer cell types and identified spatially distinct mitochondrial subpopulations with divergent metabolic states. Our results reveal previously under-appreciated spatial heterogeneity of mitochondria within cellular compartments and coordinated organization between fumarate and lipid metabolism, particularly within cellular protrusions. Our work demonstrates SPEND-hSRS as a high-resolution imaging method for multi-window, multi-organelle analysis, synchronizing molecular content to subcellular spatial organization across organelles.

### SPEND-hSRS imaging in the fingerprint window achieves sub-millimolar detection of fumarate

Fumarate stands out among TCA cycle intermediates by exhibiting several distinct vibrational features in the fingerprint region^16^. In contrast to the 1280 and 1652 cm^-1^ bands, which suffer from spectral congestion and loss of chemical specificity due to overlap with common intracellular chemical bonds, its fully symmetric COO^-^ stretch vibration at 1401 cm^-1^ is located within a spectral valley adjacent to the CH_2_ deformation mode, effectively isolating it from other dominant intracellular signals and enabling selective detection^16^.

We employed a lab-built hSRS system (**Fig. 1a left panel**; details described in **Methods**) to acquire spectra from aqueous fumarate solutions across a serial dilution series (**Fig. 1b, left panel**). Under these conditions, the LOD was determined to be 6.98 mM at a signal-to-noise ratios (SNR) of ∼3 (**Extended Data Fig. 2a, c**), with a per-pixel dwell time of 10 µs, comparable to that used in spontaneous Raman spectroscopy but achieved at substantially higher acquisition speed^16^. To further enhance sensitivity, we incorporated SPEND into the post-processing pipeline. SPEND (**Fig. 1a, middle panel**) employs a stack-permutation strategy that generates two statistically independent realizations of the same low-SNR hyperspectral dataset along the ω axis, enabling self-supervised denoising from a single acquisition, can be directly applied in batch to whole datasets acquired under identical experimental conditions for downstream analysis.

**Figure 1:**
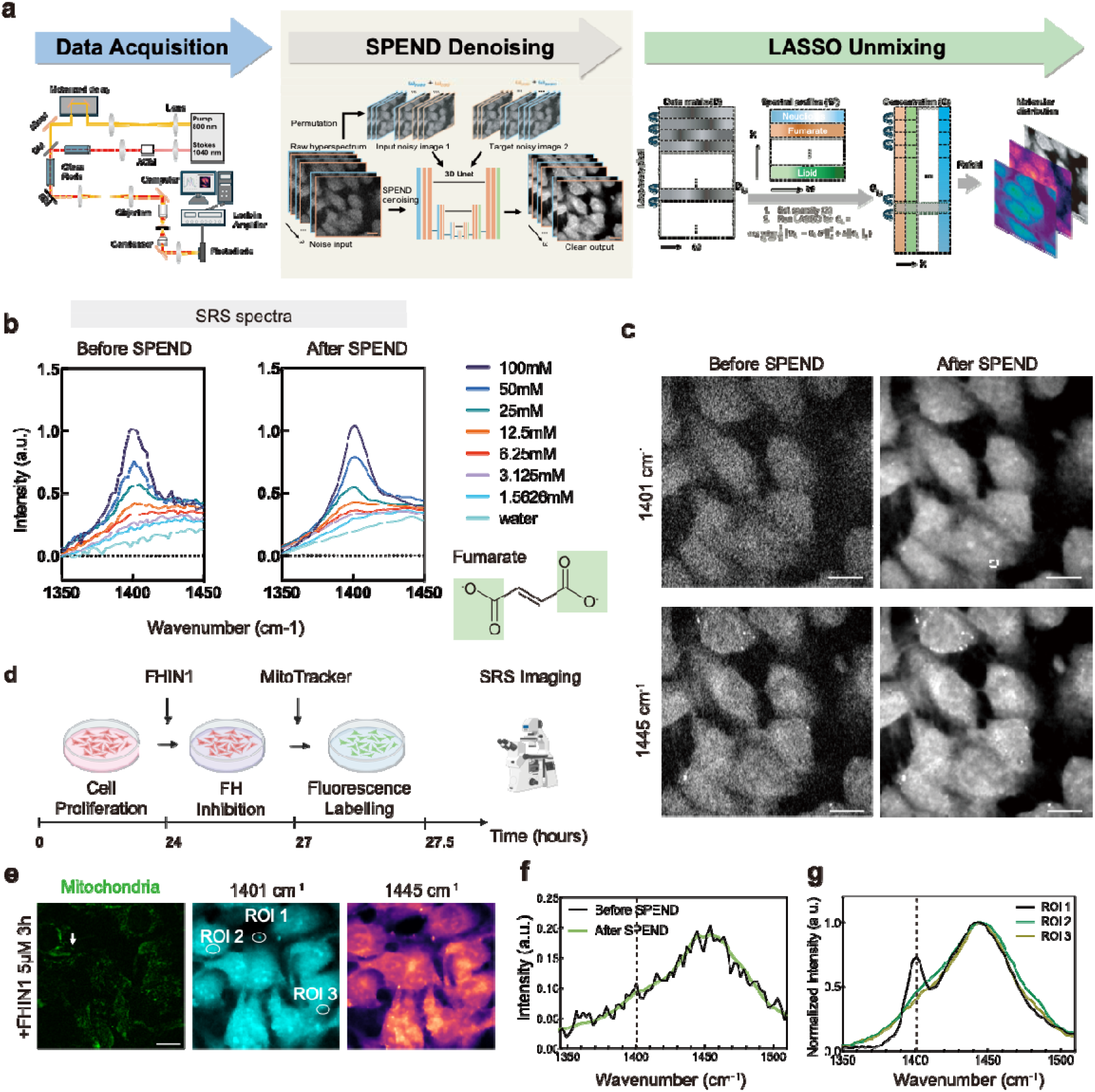
SPEND assisted NIR-SRS hyperspectral imaging in the fingerprint window reveals mitochondrial capacity for fumarate accumulation under FH inhibition. (a) Schematic of the NIR-SRS microscope and the SPEND-enabled downstream processing workflow for hyperspectral image stacks. (b) Chemical structure and spectra of fumarate aqueous solutions at different concentrations, shown before (left) and after (right) SPEND denoising. (c) Representative SRS image frames extracted from the same hyperspectral stack at 1401 cm^-1^ (fumarate) and 1445 cm^-1^ (CH_2_ deformation), shown before and after SPEND denoising. Scale bar: 15 μm. (d) Schematic of the experimental workflow combining FH inhibition (FHIN1) induced TCA-cycle blockade with hSRS imaging. (e) Representative two-photon fluorescence images of MitoTracker-labeled mitochondria (left) and corresponding SRS images at 1401 cm^-1^ (fumarate, middle) and 1445 cm^-1^ (CH_2_ deformation, right) for T24 cells under FHIN1 treatment. Scale bar: 15 μm. (f) Spectra from single mitochondrion in live T24 cell (white arrow in c) before and after SPEND denoising. (g) hSRS spectra from ROIs indicated in d.

Following SPEND denoising, noise was substantially suppressed and spectral fluctuations were markedly reduced (**Fig. 1b, right panel**), allowing us to push the LOD to 0.81 mM (**Extended Data Fig. 2b, d**). This sensitivity is well beyond the reach of spontaneous Raman spectroscopy and is fully compatible with live cell imaging without increased integration time. Applying SPEND to cellular hSRS data restored fine intracellular structures, including the nucleus, cytoplasm, and LDs, that were largely obscured in the raw hyperspectral stacks (**Fig. 2c**). These restored features showed strong spatial correspondence with the high-SNR CH-region image at 2930 cm^-1^ (**Extended Data Fig. 2e**), enabling compartment-specific spectral analysis (**Extended Data Fig. 2g**).

**Figure 2.**
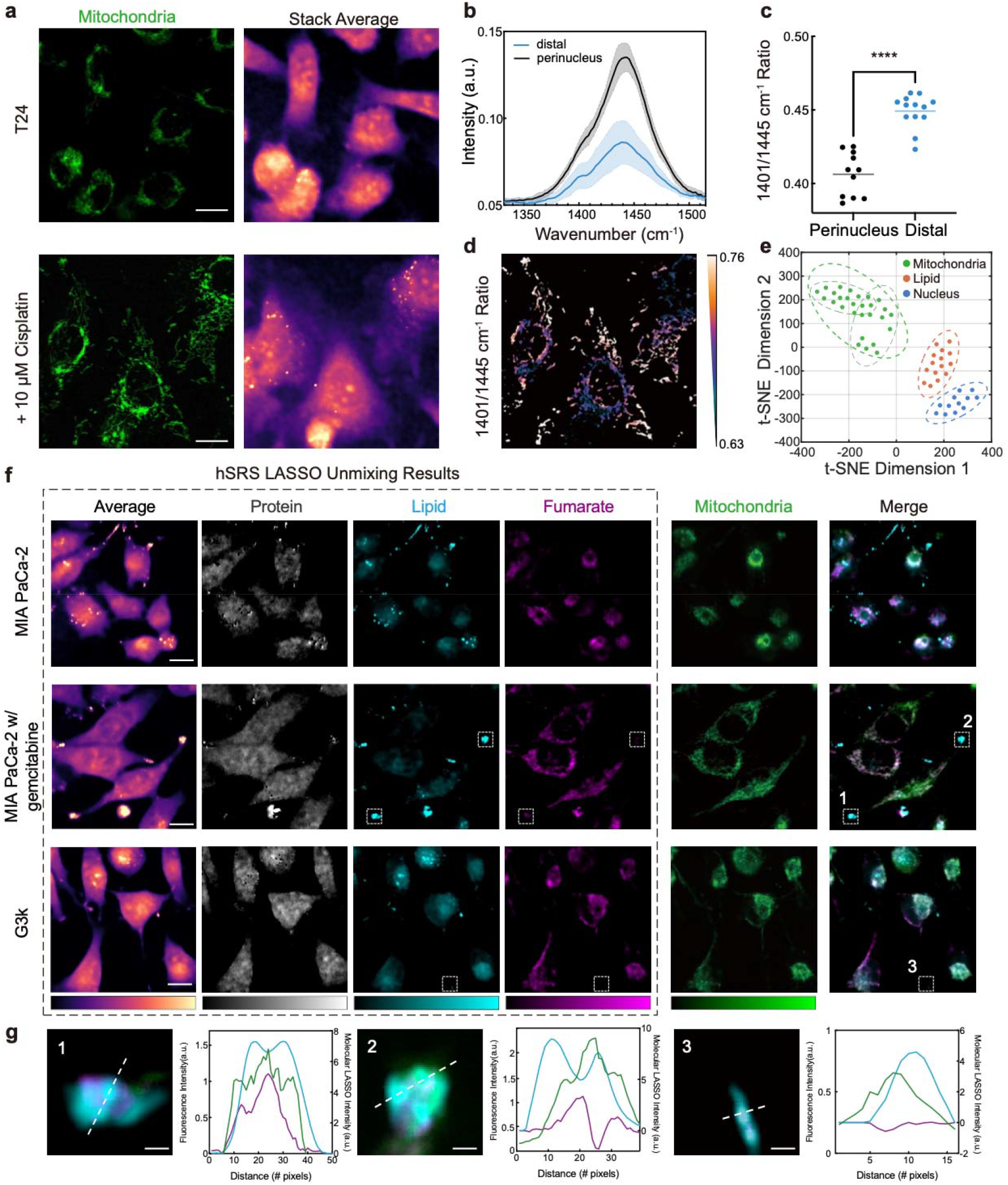
Fumarate distribution indicates mitochondria metabolic heterogeneity revealed by SPEND-SRS. (a) Two-photon fluorescence image of MitoTracker-labeled mitochondria (left) and the corresponding hSRS stack average image (right) for T24 cells. Scale bar: 15 μm. (b) hSRS spectra for different mitochondria sub-group (n=20). Graphs plus shaded areas represent mean values ± SD. (c) Quantification of the ratio of 1401 cm^-1^/1445 cm^-1^ across mitochondrial subgroups (one-way ANOVA with Tukey’s multiple-comparison test; n > 10 per group; ****P < 0.0001). (d) Color coded ratio-metric mapping for mitochondria area. Scale bar: 15 μm. (e) t-SNE spectrum dimension reduction analysis, showing clustering of subcellular compartments based on hSRS signatures. (f) Representative hSRS average, LASSO unmixing results (channels are protein, lipid, and fumarate in sequence), two-photon fluorescence for mitochondria as well as merged image of lipid, fumarate and mitochondria for MIA Paca-2 with or without gemcitabine and gemcitabine-resistant G3K cells. Each channel has the same contrast and shares a color bar. Scale bar: 15 μm. (g) Zoom-in of the dashed box in f (left) and corresponding intensity profile along the dashed line (right). Scale bar: 2.5 μm.

We first examined average spectra from different cellular compartments in fixed cells (**Extended Data Fig. 2g, left panel**), but no fumarate signal was observed at 1401 cm^-1^, which is likely attributable to chemical fixation disrupting membrane integrity, increasing membrane permeability, and thereby causing substantial loss of low-molecular-weight metabolites derived from mitochondria metabolism, in line with previous study^16,18^. In contrast, measurements in the living cells revealed a reproducible shoulder adjacent to the 1445 cm^-1^ CH_2_ deformation band (**Extended Data Fig. 2g, right panel**), a peak mainly contribute by lipid and protein. Importantly, we also observed a distinct feature near 1420 cm^-1^ arising from DNA-associated CH_2_ modes^19^, which remains spectrally separable from the fumarate signal (**Extended Data Fig. 2g, right panel**).

The substantial noise reduction achieved by SPEND further enabled spectral analysis at the level of individual mitochondrion. While spectra extracted from a single mitochondrion in fixed cells showed no detectable fumarate features (**Extended Data Fig. 2h, left panel**), spectra from a single mitochondrion in live cells exhibited a clear shoulder (**Extended Data Fig. 2h, right panel**). These observations highlight the feasibility of resolving metabolic activity among individual mitochondrion within living cells.

### TCA cycle blockade by FHIN-1 unmasks intracellular fumarate distribution

Although the achieved LOD is sufficient to detect the reported LC-MS-measured average fumarate concentration in Fumarate Hydratase (FH) deficient cells (∼9□mM)^20^, direct visualization of fumarate as a transient TCA cycle intermediate remains challenging, as its basal intracellular concentration is typically ∼10□µM or lower^21^. Under physiological conditions, mitochondrial fumarate is maintained at low steady-state levels and is therefore largely inaccessible to chemical imaging. Transient perturbation of the TCA cycle effectively converts metabolic flux into measurable fumarate accumulation, enabling spatially resolved interrogation of mitochondrial metabolism and providing a strategy that is broadly applicable across cell types.

Here, we established a FH blockade protocol (**Fig. 1d**) using a well-characterized pharmacological inhibitor, Fumarate Hydratase-IN-1 (FHIN1), interrupting downstream TCA cycle metabolism. Acute FHIN1 treatment appears to induce robust mitochondrial fumarate accumulation to levels detectable by SPEND-SRS, enabling quantitative analysis. To facilitate organelle-specific measurements, we developed a fluorescence-guided fumarate quantification pipeline (**Extended Data Fig. 3**), which allowed extraction of SRS spectra exclusively from mitochondrial regions.

Using this pipeline, we first evaluated the effect of FHIN1 dose on fumarate accumulation. No significant difference was observed between 5□µM and 20□µM FHIN1 treatment, with both conditions showing markedly elevated fumarate levels compared to untreated controls (**Extended Data Fig. 2i**). To exclude potential perturbations of mitochondrial metabolism introduced by fluorescence labeling or prolonged imaging, we varied staining duration (**Extended Data Fig. 4a**) and imaging time (**Extended Data Fig. 4c**). hSRS data acquired in both the fingerprint region, covering fumarate (1401 cm^-1^), cytochrome c (1580 cm^-1^), amide I (1655 cm^-1^), and the C–H region showed no notable spectral changes (**Extended Data Fig. 4b, d**), supporting preserved mitochondrial metabolic integrity during imaging.

We next examined fumarate accumulation in individual mitochondria in T24 bladder cancer cells (**Fig. 1e and Extended Data Fig. 2j**). FHIN-1 treatment enhanced fumarate signal intensity, rendering the characteristic spectral shoulder at 1401□cm^-1^ readily detectable (**Fig. 1f**), in contrast to untreated live cells (**Extended Data Fig. 2h**). Notably, fumarate accumulation was preferentially enriched in specific cell compartments rather than uniformly distributed, with pronounced localization observed at intercellular junctions (**Fig. 1g**), suggesting localized metabolic activity.

### Fumarate imaging indicates mitochondrial metabolic heterogeneity in cisplatin-treated cancer cells

Mitochondrial metabolism has been increasingly recognized as a central driver of cancer bioenergetics and a potential therapeutic vulnerability^22^. We therefore examined how T24 cancer cells redistribute mitochondrial organization following cisplatin treatment, a first line chemotherapy for bladder cancer. In untreated cells, mitochondria were predominantly perinuclear (**Fig. 2a, top panel**), whereas cisplatin induced redistribution toward a more dispersed cytoplasmic pattern, accompanied by a more elongated, network-like morphology (**Fig. 2a, lower panel**). This reorganization separated mitochondria into perinuclear and distal populations, based on their distance to the nucleus. Prior work has shown mitochondrial dynamics significantly affect the shape, size and position of mitochondria to support distinct cellular functions^23^. Given that functional mitochondrial heterogeneity in cancer remains incompletely understood, we asked whether these spatially defined mitochondrial populations exhibit distinct metabolic signatures.

We quantified SPEND-hSRS spectra from perinuclear and distal mitochondria. Perinuclear mitochondria exhibited higher signal near 1445 cm^-1^ (CH_2_ deformation; **Fig. 2b**), consistent with enrichment in lipid- and protein-associated content, potentially reflecting proximity to endoplasmic reticulum (ER) associated biosynthetic processes. In contrast, distal mitochondria displayed a more prominent fumarate shoulder at 1401 cm^-1^ (**Fig. 2b**). To eliminate the difference brought from concomitant changes in the CH_2_ bond, we normalized spectra to 1445 cm^-1^ and found that the distal population retained higher fumarate signal than the perinuclear population (**Extended Data Fig. 5a**).

We then quantified the ratio of 1401 cm^-1^/1445 cm^-1^ (**Fig. 2c**) and further confirmed elevated fumarate relative to CH_2_ in distal mitochondria, consistent with higher fumarate accumulation and increased TCA-linked metabolic activity in this subpopulation. Ratiometric maps further corroborated this spatial organization, revealing a gradient of mitochondrial heterogeneity from the perinuclear region toward the cell periphery (**Fig. 2d**). To further quantify absolute levels, the average fumarate concentrations were estimated by Gaussian peak fitting with linear calibration, yielding 9.60 mM in the perinuclear group and 2.81 mM in the distal group. Moreover, mitochondrial distance from the nuclear mass center correlated positively with the 1401 cm^-1^/1445 cm^-1^ ratio (**Extended Data Fig. 5b**), as supported by both Pearson and Spearman analyses (Pearson r = 0.6429; Spearman r = 0.6027). Similarly, ratiometric analysis in gemcitabine-treated MIA PaCa-2 cells also revealed different mitochondrial metabolic preference with distinct 1401 cm^-1^/1445 cm^-1^ ratios (**Extended Data Fig. 6**). Together, these data suggest the presence of metabolically distinct mitochondrial subpopulations, with one subset of mitochondria exhibiting higher fumarate and increased TCA-linked oxidative activity, and another subset, enriched in the perinuclear region, displaying a relative bias toward reduction and biosynthesis. Such mitochondrial metabolic heterogeneity may facilitate stress adaptation and survival of cancer cells^5^.

Moreover, SPEND-SRS revealed a distinct fumarate signal within these intercellular protrusions (**Extended Data Fig. 7b, c**), which colocalized with MitoTracker (**Extended Data Fig. 7b, left panel**) and was associated with LDs (**Extended Data Fig. 7a**). These observations suggest localized metabolic activity within the intercellular bridges and raise the possibility that they may participate in intercellular mitochondrial communication, consistent with previous reports of mitochondrial transfer through tunneling nanotubes (TNTs)-like structures^24,25^.

As noted above, the fumarate COO^-^ bond at 1401 cm^-1^ lies in a spectral valley adjacent to the CH_2_ deformation mode, raising the possibility of resolving multiple chemical components within this information-rich window from a single hyperspectral acquisition. We therefore first validated organelle-level spectral separability among mitochondria, LDs, and nucleolus. t-SNE analysis of organelle spectra showed clear clustering by compartment (**Fig. 2e**). Interestingly, mitochondria further separated into subclusters, consistent with the mitochondrial heterogeneity inferred above. We then performed pixel-wise LASSO unmixing on SPEND-hSRS image stacks (**Fig. 1a, right panel**; details described in **Methods**), generating metabolite maps of key biochemical components including fumarate, protein and lipid (**Fig. 2f**) by using the standard spectra (**Extended Data Fig. 5e**). We further analyzed cells without FHIN1 treatment (**Extended Data Fig. 8**), the inferred fumarate signal was weak and largely indistinguishable from background, making reliable and specific spatial localization impossible. Notably, the fumarate spatial distribution was not fully captured by MitoTracker labeling (**Fig. 2f**). This discrepancy likely reflects the presence of fumarate beyond mitochondria, whereas MitoTracker selectively labels mitochondria.

Cancer-cell protrusions are widely considered part of an epithelial-to-mesenchymal (EMT)-like transition toward a more invasive phenotype and have been reported across many solid tumors^26^. EMT is associated with remodeling of the actin cytoskeleton, which is essential to organelle transport and dynamics^27^. We then asked whether mitochondrial metabolism is spatially reprogrammed during stress-associated morphological remodeling. We treated MIA PaCa-2 cells with gemcitabine and observed a marked shift toward a spindle-like morphology with redistribution of lipid droplets into protrusions (**Fig. 2f**). While MitoTracker signal remained enriched in perinuclear regions, fumarate signal was strongly elevated within protrusions along with lipid signal (**Fig. 2g**, zoom in figure 1 and 2). Estimated fumarate concentrations were 7.23 mM in the perinuclear group and 2.28 mM in the distal group. Moreover, the fumarate distribution in dashed box 1 was broader than in dashed box 2, consistent with previously reported patterns of polarized metabolic organization during the transition^28^. By contrast, this coupling was diminished in G3k cells (**Fig. 2f**), a MIA PaCa-2 derived gemcitabine-resistant line. Although G3k exhibited similar morphological features (**Extended Data Fig. 5c**), but the lipid and MitoTracker signal in the protrusion was no longer associated with fumarate (**Fig. 2g**, zoom in figure 3), which may reflect metabolic adaptation to prolonged chemotherapy stress. Finally, single-cell integrated fumarate intensity quantification showed the highest intracellular fumarate levels in gemcitabine-treated MIA PaCa-2 cells, with reduced levels in G3k cells (**Extended Data Fig. 5f**) and no difference between MIA PaCa-2 and G3k, supporting a transition from acute stress–associated response to an adapted mitochondria metabolic state.

**Figure 3.**
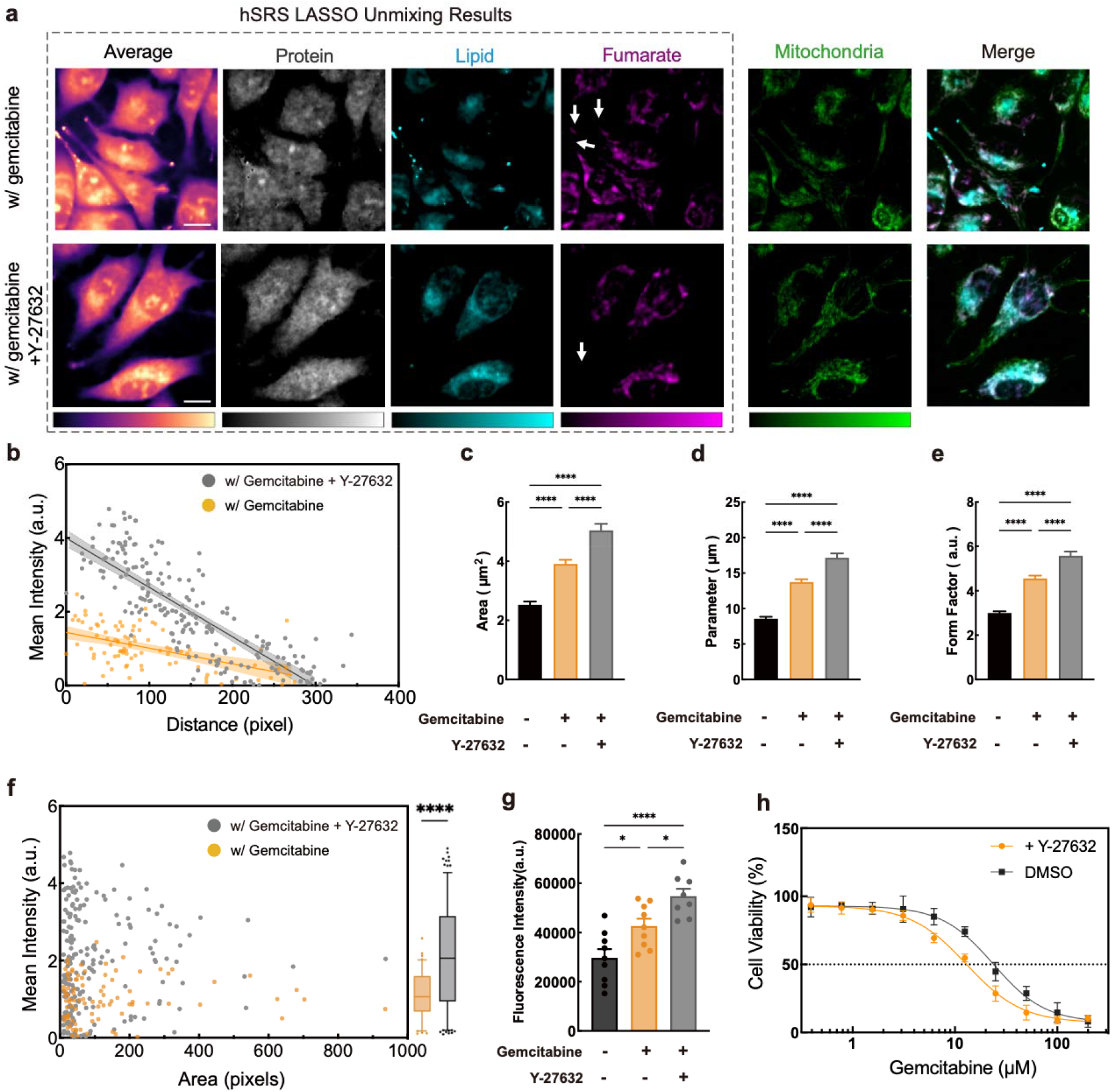
Fumarate imaging reveals mitochondrial metabolic changes upon pharmacological perturbation in Rho pathway. (a) Representative hSRS average, LASSO unmixing results for hSRS images (channels are protein, lipid, and fumarate in sequence.), two-photon fluorescence of MitoTracker-labeled mitochondria, and a merged image of lipid, fumarate, and mitochondria from MIA PaCa-2 cells treated with gemcitabine with or without a ROCK inhibitor. Scale bar: 15 μm. (b) Relationship between mean fumarate intensity per mitochondrion and distance to the nuclear mass center in gemcitabine-treated MIA PaCa-2 cells without (n = 91) or with (n = 197) ROCK inhibitor. Solid line, linear regression; shaded band, 95% confidence interval. (c–e) Quantification of mitochondrial morphology under the indicated gemcitabine and ROCK inhibitor conditions: (c) area, (d) parameter and (e) form factor. Bars show mean ± SEM. (f) Association between mitochondrial mean fumarate intensity and mitochondrial area in gemcitabine-treated MIA PaCa-2 cells without (n = 91) or with (n = 197) ROCK inhibitor. Boxplots summarize the distribution of mitochondrial fumarate signal before and after ROCK inhibition; center line, median; box, interquartile range; whiskers, 5th–95th percentiles. (g) Spectrophotometric fluorescence intensity measurement of intracellular ROS levels in MIA PaCa-2 cells under the indicated drug treatments. For (c–e) and (g), statistical significance was assessed by one-way ANOVA with Tukey’s multiple-comparison test (*P < 0.05, ****P < 0.0001). (h) Gemcitabine concentration-dependent MIA PaCa-2 cell viability with and without Y-27632 treatment, the bars indicate means ± SEM.

### Pharmacological disruption of Rho pathway induces mitochondrial metabolic shifts

Because the Rho pathway plays a key role in spatiotemporal coordination of cytoskeletal dynamics that drives protrusion formation during cell migration^29^, we asked whether pharmacological regulation of Rho signaling could rewire mitochondrial metabolism. We co-treated MIA PaCa-2 cells with gemcitabine and the Rho-kinase (ROCK) inhibitor Y-27632. Surprisingly, we observed a significant remodeling of fumarate distribution in LASSO unmixing results, despite minimal discernible differences in the MitoTracker channel (**Fig. 3a**). Notably, while under ROCK inhibition, cells continued to transport LDs toward protrusive ends (**Fig. 3a, bottom panel; Extended Data Fig. 9a**), fumarate became preferentially concentrated higher in the perinuclear region rather than in distal protrusions, indicating a spatial remodeling of mitochondrial metabolic activity. Quantifying each mitochondrion by its distance to the nuclear mass center revealed distinct distance–fumarate relationships between conditions (**Fig. 3b**): the fumarate signal decayed more rapidly with distance in the gemcitabine and ROCK inhibition group than with gemcitabine alone (k = −0.0134 vs −0.0044), an effect that was even clearer after normalization (**Extended Data Fig. 9b**). In contrast, gemcitabine-only cells exhibited markedly greater variance in fumarate at larger distances, suggesting enhanced distal mitochondrial heterogeneity under chemotherapy stress, potentially reflecting impaired redistribution of lipid droplets and mitochondria and their retention in the perinuclear region.

In parallel, ROCK inhibition subtly altered mitochondrial morphology (**Fig. 3a, Extended Data Fig. 9c**). Given that mitochondrial size and shape report on organelle dynamics and functional state^23^, we quantified area, perimeter, and form factor and observed a concerted shift under Y-27632 and gemcitabine co-treatment toward expanded, more elongated and structurally complex mitochondria (**Fig. 3c–e**). The relationship between mitochondrial area and average fumarate intensity revealed an enriched subgroup in the ROCK inhibition group characterized by enlarged morphology and elevated fumarate levels (**Fig. 3f**), indicating a metabolically active state. Since reactive oxygen species (ROS) can function as adaptive signals for mitochondria under stress conditions, we measured intracellular ROS levels and found that the gemcitabine and Y-27632 co-treatment group exhibited the highest ROS levels (**Fig. 3g**), consistent with metabolic and morphological changes. ROS is often regarded as simple by-products of respiration and associated with dysfunction under normoxia, ROS levels can also report changes in respiratory flux and accompany stress-adaptive mitochondrial remodeling^30^. These data collectively support a model in which ROCK inhibition not only remodels cytoskeletal organization but also alters the spatial distribution of mitochondrial metabolism. It redistributes TCA cycle activity to perinuclear compartments and amplifies a distinct, TCA cycle-active mitochondrial subpopulation during chemotherapy. Consistently, our cell viability assays revealed that co-treatment with the ROCK inhibitor significantly reduced the IC50 of gemcitabine in MIA PaCa-2 cells (**Fig. 3h**), indicate the importance of LD and mitochondria interaction in the survival mechanism for cancer cells under chemotherapy stress.

### C-H SRS imaging shows intracellular interplay between lipid metabolism and mitochondrial metabolism

Given that we simultaneously observed LD accumulation and altered mitochondrial metabolic activity in cell protrusions under different drug treatments, we asked whether mitochondrial metabolic shifts are coupled to LD content. LD localization and composition are dynamically regulated^31^, and LD can functionally interact with mitochondria as an energy reservoir. To test whether perturbing mitochondrial metabolism influences LD composition in protrusions, we acquired hSRS images in the C–H window to probe LD chemical composition.

Consistent with the fingerprint-region analysis, we observed LD accumulation in pharmaceutically perturbed MIA PaCa-2 cells (**Fig. 4a**) and in G3k cells (**Extended Data Fig. 10a**). We compared LD spectra in protrusions before and after ROCK inhibition (**Extended Data Fig. 10b**). The LD in gemcitabine-treated MIA PaCa-2 cells exhibited a distinct peak at 2870 cm^-1^, a feature associated with cholesterol. Moreover, we observed ROCK inhibition further increased the overall LD signal intensity. These results suggest that fatty acids stored in LD at protrusions may normally be mobilized to fuel mitochondrial energy production, whereas ROCK inhibition impairs this utilization (**Fig. 4b**).

**Figure 4:**
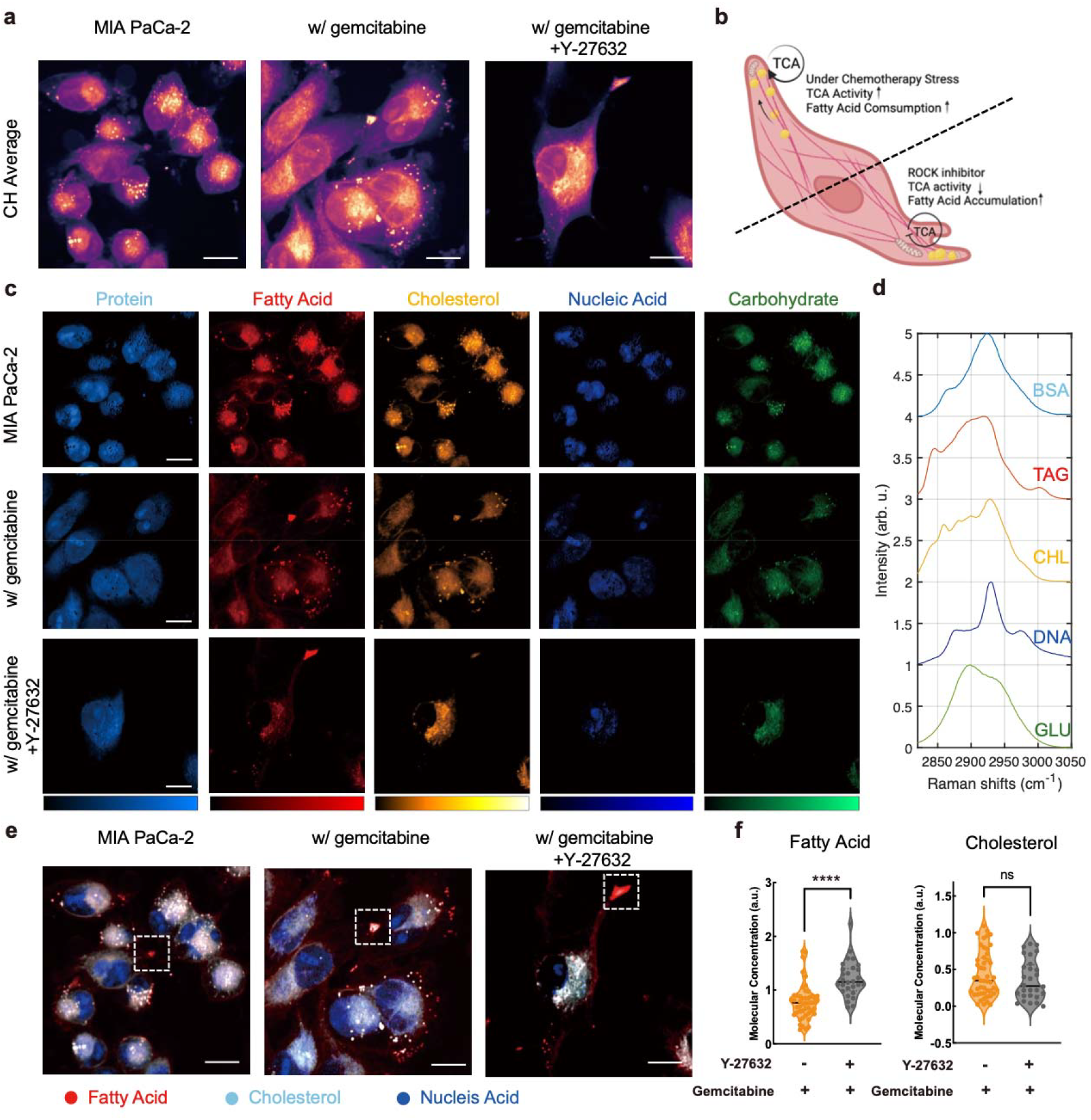
C-H SRS imaging reveals interplay between lipid metabolism and mitochondria metabolism. (a) Representative hSRS average for MIA PaCa-2, MIA PaCa-2 treat with gemcitabine, MIA PaCa-2 treat with gemcitabine and ROCK inhibitor. Scale bar: 15 μm. (b) Schematic illustrating chemotherapy-induced LDs accumulation for energy supply in cell protrusions and its perturbation by ROCK inhibition. (c) LASSO unmixing results for hSRS images (channels are fatty acid, cholesterol, nucleic acid and carbohydrate in sequence). Scale bar: 15 μm. (d) Normalized SRS reference spectra of the target metabolites for LASSO spectral unmixing. Protein, fatty acid, cholesterol, nucleic acid and carbohydrate are represented by bovine serum albumin (BSA), triglyceride (TAG), cholesterol (CHL) powder, glucose (GLU) and cell-extracted deoxyribonucleic acid (DNA), respectively. Spectra are vertically offset for clarity. (e) Merged image of TAG, cholesterol and Nucleic Acid from LASSO results. Scale bar: 15 μm. (f) Individual LDs molecular concentration quantification of TAG and cholesterol in gemcitabine-stressed MIA PaCa-2 cells with or without ROCK inhibitor (n > 30). Statistical significance was assessed by two-tailed Student’s t-test (ns, not significant; ****P < 0.0001).

To identify the lipid species contributing to LDs, we applied the LASSO spectral unmixing in the CH window to separate different compositions including protein, fatty acid, cholesterol, carbohydrate and nucleic acid (**Fig. 4c**) using standard references (**Fig. 4d**). Under gemcitabine treatment, LDs in MIA PaCa-2 protrusions showed an increased cholesterol fraction (**Fig. 4e**) relative to untreated MIA PaCa-2 cells. After ROCK inhibition, LDs accumulated more prominently in protrusions and triglycerides became the dominant species again (**Extended Data Fig. 10e**). This trend is consistent with our fingerprint-region findings.

In gemcitabine-treated MIA PaCa-2 cells, mitochondria in protrusions showed strong fumarate signal (**Fig. 2f**), which disappeared upon Y-27632 treatment (**Fig. 3a**), suggesting that ROCK inhibition impedes the transfer or utilization of lipid-derived metabolic substrates from LDs to mitochondria. In this scenario, triglycerides accumulate within LDs, are less efficiently consumed, and consequently form larger droplets (**Extended Data Fig. 10e**). We further quantified the composition of individual LDs (**Fig. 4f**). When cholesterol levels across individual LDs were comparable, cells treated with Y-27632 exhibited higher triglyceride content, further supporting impaired fatty-acid oxidation under ROCK inhibition. Our data suggest that the ROCK pathway can interfere with LDs–mitochondria coupling and impair lipid supply, although the underlying molecular mechanisms remain to be elucidated. Taken together, LDs accumulation in protrusions may represent an adaptive strategy that helps cancer cells survive under stress.

## Discussion

Historically, mitochondrial dysfunction has been viewed as a hallmark of cancer cells^32^. Emerging research suggests that individual cells harbor discrete mitochondrial subpopulations, each possessing specialized metabolic functions tailored to specific cellular preferences^5^. It is recognized that mitochondrial structure and function are dynamically tuned to local demands for energy production, biosynthesis, repair, and renewal^2,4^. However, directly interrogating mitochondrial metabolism in situ at the sub-organelle level has remained technically challenging.

Here, we introduce SPEND-hSRS imaging as an approach to probe mitochondrial metabolic organization using fumarate as an endogenous readout. SPEND denoising of hSRS data in the fingerprint region enables quantitative analysis for the low-abundance biomolecules. In this study, SPEND improved the LOD for fumarate in aqueous standards to 0.81 mM, which is ∼10 times better than spontaneous Raman microscopy^16^. Paired with the FH blockade strategy, this workflow is broadly applicable across cell types without added optical complexity or genetic manipulation.

We first observed chemotherapy induced mitochondrial dispersion in cancer cells. Ratiometric analysis revealed spatially patterned fumarate enrichment despite apparently uniform MitoTracker labeling, underscoring the metabolic heterogeneity information provided by SPEND-SRS beyond conventional fluorescence labeling. We then leveraged pixel-wise LASSO spectral unmixing to reconstruct intracellular fumarate maps, which uncovered an association between fumarate signal and LDs within protrusive regions following gemcitabine treatment. To test whether this organization was coupled to protrusion dynamics, we inhibited ROCK signaling and observed coordinated changes that linked mitochondrial morphological features with fumarate abundance, suggesting mitochondria morphology has strong correlation with TCA metabolism.

Extending the analysis to the C-H window further suggested that mitochondria metabolic states are accompanied by lipid turnover: gemcitabine-treated MIA PaCa-2 cells showed LDs accumulation with reduced triglycerides, this pattern is consistent with triglycerides and can be used to support mitochondrial β-oxidation^33^. By contrast, ROCK inhibition reduces this lipid utilization, leading to triglyceride storage in LDs. We demonstrate that MIA PaCa-2 cells dynamically reorganize mitochondria-LDs interactions under chemotherapeutic stress to support stress adaptation. Disrupting this inter-organelle coupling enhances cellular sensitivity to chemotherapy, highlighting it as a potential therapeutic vulnerability. In parallel, combining SPEND-hSRS imaging with immunofluorescence labeling of mitochondria- and LD-associated proteins^34^ would help futher understanding the spatial relationships observed here and provide mechanistic insight into the molecular basis of inter-organelle interaction.

The SPEND-hSRS technique expands chemical imaging access to fingerprint window that was previously obscured by noise and difficult to utilize for resolving organelle composition. A remaining limitation is spectral bandwidth. Recently developed super-broadband stimulated Raman scattering spectroscopy can achieve 100-fold increase in speed than spontaneous Raman^35^. Further development of broader-bandwidth and higher-sensitivity coherent Raman microscopy implementations could further facilitate organelle-level metabolic imaging.

Finally, fumarate is more than a convenient reporter of TCA-associated metabolism. As an oncometabolite, its accumulation can actively reshape cellular physiology, influencing processes such as mitophagy, redox stress, and tumor progression^36,37^. In this sense, SPEND-hSRS can be used to investigate how fumarate contributes to these pathological processes. Taken together, our results establish SPEND-hSRS as a practical platform for mapping fumarate at subcellular resolution and for revealing mitochondrial heterogeneity in living cells. By linking organelle-scale metabolic organization to protrusive behavior and lipid turnover, this approach provides insight into cancer cell invasive behavior and suggests potential therapeutic vulnerabilities.

## Methods

### Hyperspectral SRS and two-photon excitation fluorescence (TPEF) microscope

A lab-built hyperspectral stimulate Raman scattering microscope (**Fig. 1a**), previously published^38^, is used to perform hyperspectral SRS imaging. A femtosecond pulse laser (Insight, DeepSee+, spectra-Physics) operating at 80MHz with two synchronized beams, a tunable pump beam ranging from 680nm to 1300nm, a fixed stokes beam at 1040nm. The pump beam is tuned to 800nm for the C-H region, 907nm for the fingerprint 1300 - 1500 cm^-1^ region. The Stokes beam is modulated at 2.5MHz by an acoustic optical modulator (1205-C, Isomet) and chirped by a 15cm glass rod (SF57, Schott) before the merging of two beams. The combined two beams were chirped by five glass rods to picosecond pulse. A motorized linear stage is used to tune the time delay between the pump and Stokes pulse which corresponds to the Raman shift of chemical bonds. A 2D Galvo scanner (GVS102, Thorlabs) is used for laser scanning. The combined beam is sent to the sample through a 60X water immersion objective (NA=1.2, UPlanApo/IR, Olympus). After interacting with the sample, the beam is collected by an oil condenser (NA=1.4, U-AAC, Olympus). A photodiode (S3994-01, Hamamatsu) is used to collect signals after filtering the Stokes beam. The lock-in amplifier (UHFLI, Zurich Instrument) is used to extract high frequency signals.

The TPEF imaging was performed on the same SRS laser scanning microscopy platform and detected by a photomultiplier tube (PMT) (H422-40, Hamamatsu Photonics). After the PMT, a pre-amplifier was used to amplify the signal before sending the signal to the data acquisition card (PCIe-6363, National Instruments).

### Self-supervised denoising of hyperspectral SRS images

We implemented the Self-Permutation Noise2Noise Denoiser (SPEND) using a four-layer U-Net architecture, implemented with Tensorflow^15^. The network was trained on three independent hyperspectral datasets. Each stack contains 600 * 600 pixels and 100 frames. To improve robustness and reduce overfitting, augmentation was applied to each dataset using rotations and flips, resulting in a four-fold expansion per dataset and a total of 12 stacks used for training. Augmentation preserves the physical statistics of the noise while increasing the diversity of spatial structures encountered during training. Training and inference were conducted on an NVIDIA RTX 4090 GPU.

### Pixel-wise least absolute shrinkage and selection operator (LASSO) spectral unmixing

Raw hyperspectral SRS images were acquired as 3D data stacks, in which each spatial pixel contains a full SRS spectrum. To convert these hyperspectral datasets into metabolite concentration maps, we performed pixel-wise spectral unmixing using LASSO algorithm.

For unmixing in the fingerprint region, reference spectra were obtained from 500 mM fumarate in aqueous solution and triglyceride standards; a nucleolus reference spectrum was extracted from compartment-averaged cellular spectra. These references represent fumarate, lipids, and nucleolus, respectively. For unmixing in the C-H stretching region, reference spectra of bovine serum albumin (BSA), triglyceride (TAG), cholesterol (CHL), glucose (GLU), and ribonucleic acid (RNA) were measured and used to represent proteins, lipids, cholesterol, carbohydrates, and nucleic acids, respectively.

To mitigate spectral crosstalk arising from overlapping Raman bands, we used an L1-norm sparsity regularization on the per-pixel concentration vector. Implementation details and validation of the algorithm have been described previously^44^.

### Cell culture and imaging preparation

MIA PaCa-2 (CRL-1420) and T24 (HTB-4) cells were obtained from the American Type Culture Collection (ATCC). Gemcitabine-resistant G3K cells were derived from parental MIA PaCa-2 cells through repeated exposure to gemcitabine^39^. All cell lines were authenticated and routinely confirmed to be mycoplasma-free using the MycoAlert assay (Lonza, LT07-318).

Cells were cultured in high-glucose Dulbecco’s modified Eagle’s medium (DMEM, Gibco) supplemented with 10% fetal bovine serum (FBS, Gibco) and penicillin-streptomycin (P/S, 100 U mL^-1^; Gibco) and maintained at 37□°C in a humidified incubator with 5% CO_2_.

For imaging experiments, cells were seeded in 35-mm glass-bottom dishes. For live-cell imaging, the culture medium was replaced with Live Cell Imaging Solution (Invitrogen, A59688DJ) after three washes with PBS. For fixed-cell imaging, cells were fixed with 4% paraformaldehyde (PFA) in PBS (Boston BioProducts, BM-155), washed three times with PBS, and then stored in PBS for imaging.

### FHIN1 treatment

Fumarate hydratase-IN-1 (FHIN1; MedChemExpress, HY-100004) was used to induce acute fumarate accumulation prior to SRS imaging. FHIN1 was prepared as a 10 mM stock solution in DMSO and diluted to a final working concentration of 5 μM in complete culture medium. Cells were treated with FHIN1 for 2 h, washed with PBS, and then subjected to mitochondrial staining.

### Cell stress treatment

To establish a chemotherapy stress model, T24 cells were treated with 10 μM cisplatin (Sigma, 232120) and MIA PaCa-2 cells were treated with 10 μM gemcitabine (SelleckChem, S1714) for 24 h prior to SRS imaging.

To perturb Rho pathway signaling, the ROCK inhibitor Y-27632 (SelleckChem, S1049) was used. Y-27632 was dissolved in DMSO to generate a 10 mM stock and subsequently diluted in complete culture medium to 10 μM. Cells were treated with Y-27632 for 24 h prior to imaging.

### Mitochondria fluorescence staining

Live-cell mitochondria were stained with MitoTracker™ Green (Thermo Fisher Scientific, M7514). MitoTracker Green fluorescence was acquired using two-photon excited fluorescence (TPEF) sequentially with SRS imaging (excitation: 950 nm; emission collected on a PMT through a 520/30 nm bandpass filter). For fixed-cell mitochondria morphological analysis, mitochondria were labeled with MitoBrilliant™ 646 (Tocris, 7700) and imaged on an Olympus FV3000 confocal microscope using a UPlanSApo 60× water-immersion objective.

### ROS measurement

Intracellular reactive oxygen species (ROS) were quantified using the DCFDA assay (Abcam, #ab113851). Adherent cells were seeded in black, clear-bottom 96-well plates (25,000 cells per well) and allowed to attach overnight. Corresponding drug was given at the same time. Cells were incubated with DCFDA (20 μM) for 45 min at 37 °C in the dark, followed by removal of the dye solution and replacement with buffer or phenol red–free complete medium containing 10% FBS. Fluorescence was measured immediately in endpoint mode using a microplate reader (Ex/Em = 485/535 nm).

### Cell viability assay

MTS assay (Abcam, #ab197010) has been used to measure cell viability. After overnight seeding of cells at 96-well plates at densities of 2000 cells per well, chemotherapy treatment was introduced to the cell culture at indicated concentrations for 24 h. After 1.5 h incubation with MTS reagent, cell viability was evaluated through absorbance at 490 nm measured by plate reader.

### Spontaneous Raman spectroscopy

Confocal Raman was performed by a commercial Raman microscope (LabRAM HR Evolution, Horiba) at room temperature. A 532-nm diode laser was used to excite the sample through a 40× water immersion objective (Apo LWD, 1.15 N.A., Nikon). The total data acquisition time was 15 s using the LabSpec 6 software.

### Statistical Analysis

GraphPad Prism 10 software was used for statistical analyses. For parametric data, student t-tests were used for non-paired data and paired t-test for paired data. For multiple comparisons, one-way analysis of variance (ANOVA) test was used. Error bars, *P* values and statistical tests are reported in the figures and figure legends. For correlative analysis, Pearson’s method for linear regression and Spearman’s method for the nonlinear regression were used. All images were processed in the ImageJ (https://imagej.net/ij/). LASSO spectral unmixing is performed in MATLAB 2024b. Experiments were performed as biological triplicates.

### Figure construction

Graphs were created in GraphPad Prism 10 and assembled into figures in Adobe illustrator 2025. The schematics in Fig 1d and 4b and Extended Data Fig. 4 were created using BioRender (Created in BioRender. Sun, D. (2026) https://BioRender.com/wqjtaaj).

## Acknowledgements

The authors thank Dr. Zhongyue Guo for the beneficial discussion in the project conceptualization and the graphical support. And we thank Boston University Biomedical Engineering Micro and Nano Imaging (MNI) facility for providing the confocal fluorescence microscope. This work is supported by National Institutes of Health under award Number R35GM136223 and R01EB035429.

## Author contributions

D.S. and J.X.C. conceived the study. D.S. performed experiments and analyzed the data. G.D. developed the SPEND. H.L. helped with experiments. H.L and G.C. helped with data discussion. C.W., S.B. and S.E.B. helped with project conceptualization. All authors participated in discussing and finalizing the manuscript.

## Data and Code Availability

All data related to the work are available in the article and supplementary information in this paper and are available upon reasonable request to the corresponding authors. All relevant code, including SPEND denoising and LASSO spectral unmixing code, can be accessed at the Cheng Lab GitHub page (https://github.com/buchenglab).

