## Supplementary Figure for "SPEND-hSRS imaging of fumarate uncovers mitochondrial metabolic heterogeneity"

### Extended Data Figure

Dingcheng Sun[1, 3], Guangrui Ding[2, 3], Haonan Lin[2, 3, #], Guo Chen[2, 3], Chun-Chin Wang[4, 5], Seema Bachoo[4, 5], Sarah E. Bohndiek[4, 5], Ji-Xin Cheng[1, 2, 3*]

[1]Department of Biomedical Engineering, Boston University, Boston, MA, USA, 02215

[2]Department of Electrical and Computer Engineering, Boston University, Boston, MA, USA, 02215

[3]Photonics Center, Boston University, Boston, MA, USA, 02215

[4]Department of Physics, University of Cambridge, JJ Thomson Avenue, Cambridge, CB3 0HE, UK

[5]Cancer Research UK Cambridge Institute, Robinson Way, Cambridge, CB2 0RE, UK

[#] Current address: The Wallace H. Coulter Department of Biomedical Engineering, Georgia Institute of Technology and Emory University, Atlanta, GA, 30332

a

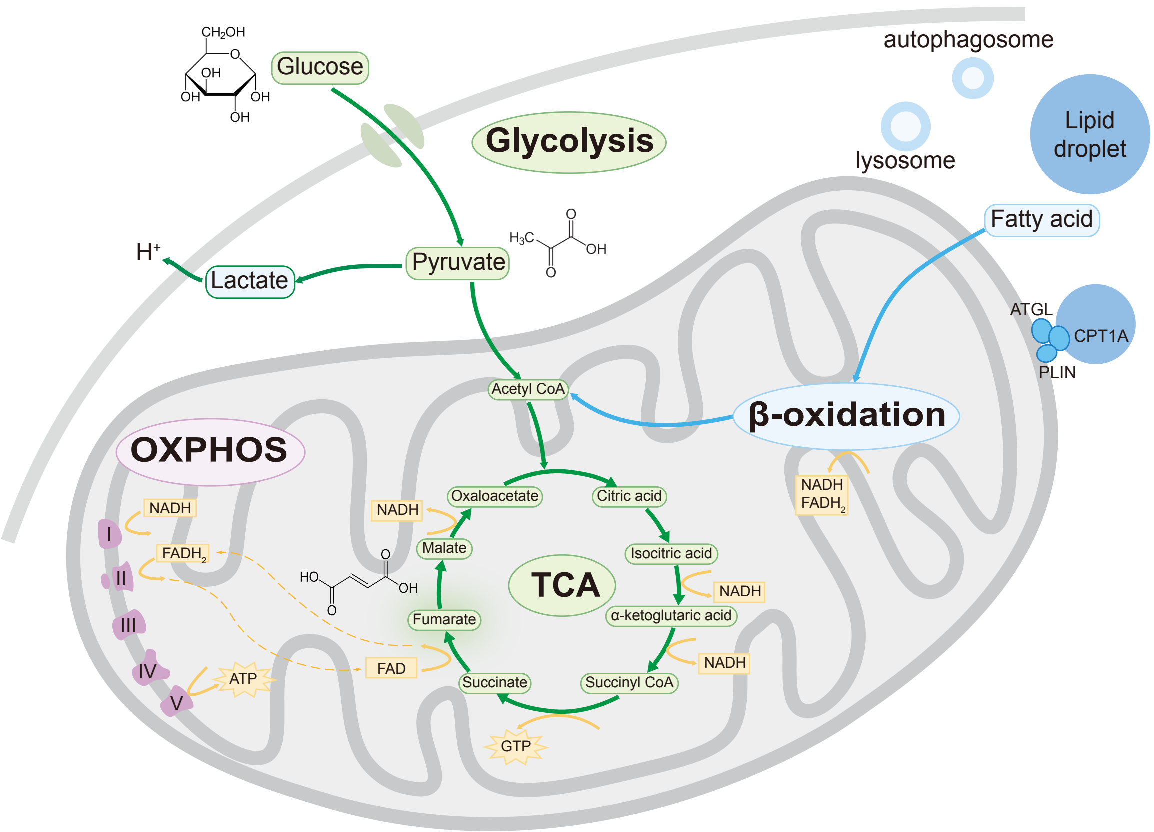


b

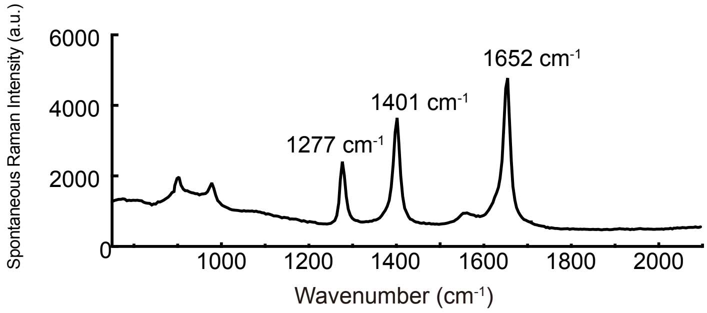


**Extended Data Figure 1: Fumarate locate in the TCA cycle displays unique Raman spectral features.**

(a) Overview of mitochondrial energy metabolism. Glucose is metabolized by glycolysis to generate pyruvate, which enters mitochondria and is converted to acetyl-CoA. Acetyl-CoA fuels the tricarboxylic acid (TCA) cycle, producing reducing equivalents (NADH and $FADH_2$) and GTP. In parallel, fatty acids released from lipid droplets undergo mitochondrial β-oxidation to generate acetyl-CoA as well as NADH and $FADH_2$. NADH and $FADH_2$ donate electrons to the respiratory chain to drive oxidative phosphorylation (OXPHOS) and ATP production. (b) Spontaneous Raman spectra for 500 mM fumarate aqueous solution.

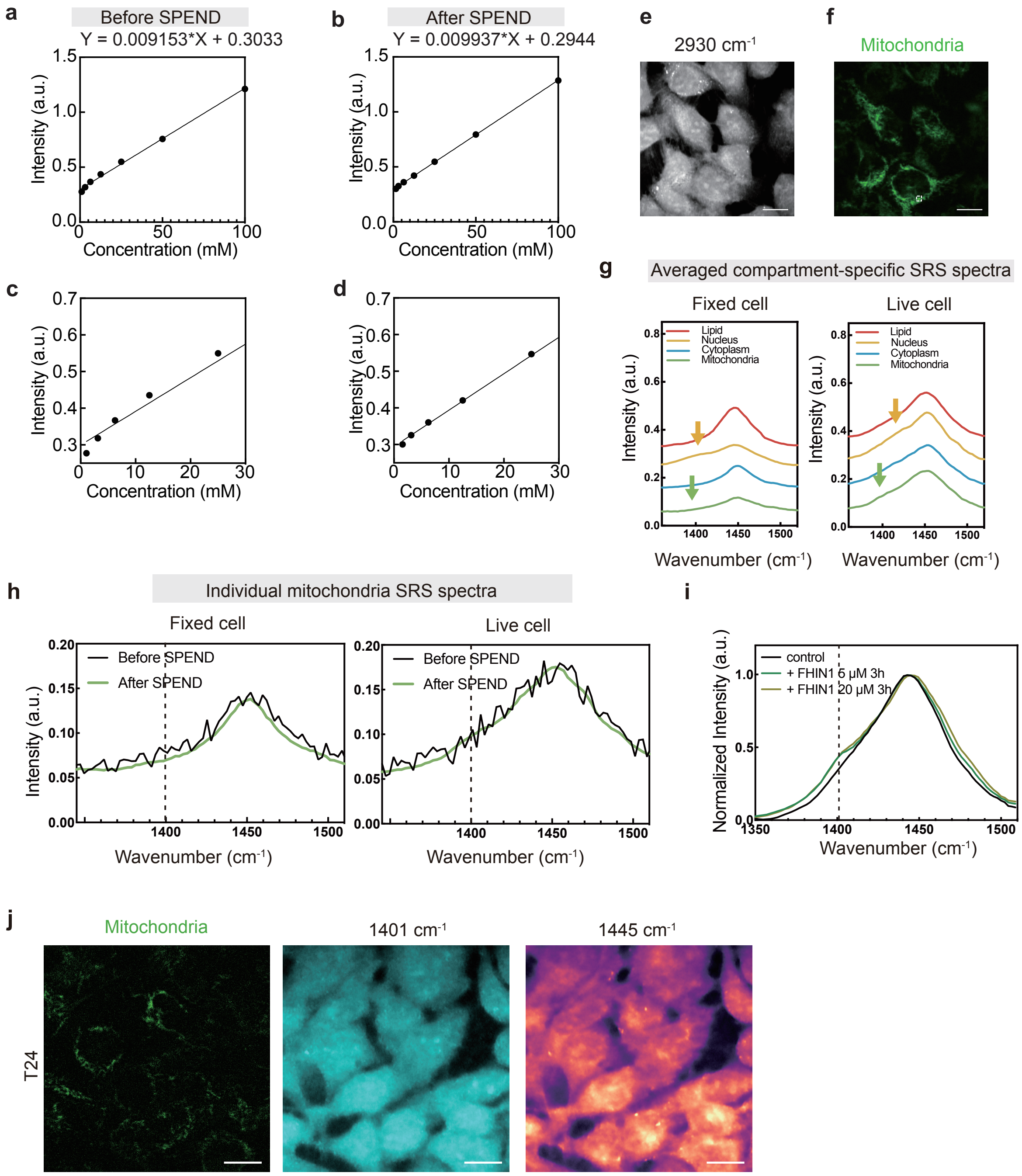


**Extended Data Figure 2: Improved fumarate detection sensitivity by SPEND-SRS imaging.**

(a, b) Limit of detection (LOD) analysis for the fumarate at 1401 $cm^{-1}$ before (a) and after SPEND (b). SRS intensity is plotted as a function of fumarate concentration with linear regression (fit equation shown). (c, d) Zoom in for the linear regression for smaller range of concentrations in a and b. The estimated LOD decreases from 6.98 mM before SPEND to 0.81 mM after SPEND denoising. (e) Representative live-cell SRS image in the C-H window acquired together with (f) MitoTracker labeling (corresponding to Fig. 1c). Scale bar: 15 μm. (g) Averaged compartment-specific spectra (n > 12 per compartment). Spectra are vertically offset 0.1 for clarity and color coded according to the compartment. (h) Representative spectrum from a single-mitochondrion region before and after SPEND processing, illustrating improved spectral fidelity under low-SNR conditions. (i) Average mitochondrial SRS spectra measured after different FHIN1 treatment dosages. (j) Representative two-photon fluorescence images of MitoTracker-labeled mitochondria (left) and corresponding SRS images at 1401 $cm^{-1}$ (fumarate, middle) and 1445 $cm^{-1}$ ($CH_2$ deformation, right) for untreated T24 cells. Scale bar: 15 μm.

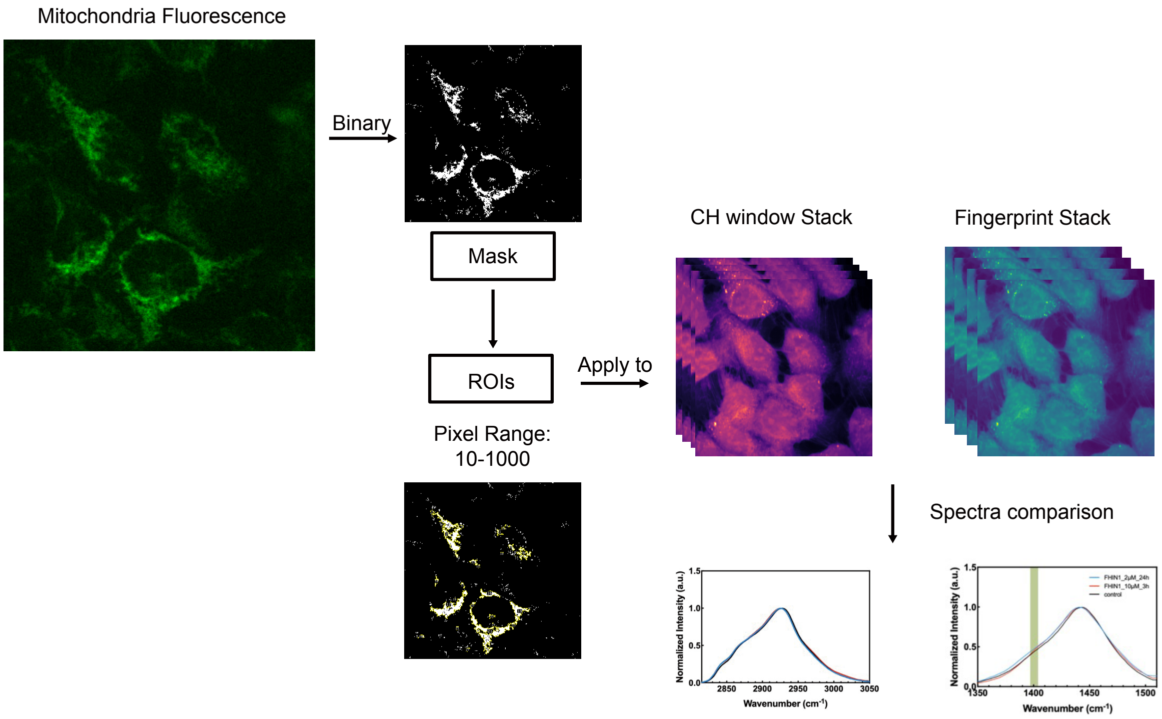


**Extended Data Figure 3: MitoTracker-guided quantification of mitochondrial fumarate signals.**

Fluorescence-guided workflow for extracting mitochondria-specific hSRS spectra. MitoTracker fluorescence images were binarized to generate a mitochondrial mask, which was used to define ROIs. After pixel filtering (10–1000), the ROIs were applied to both CH and multiple fingerprint window to extract mitochondria-specific spectra for subsequent comparison.

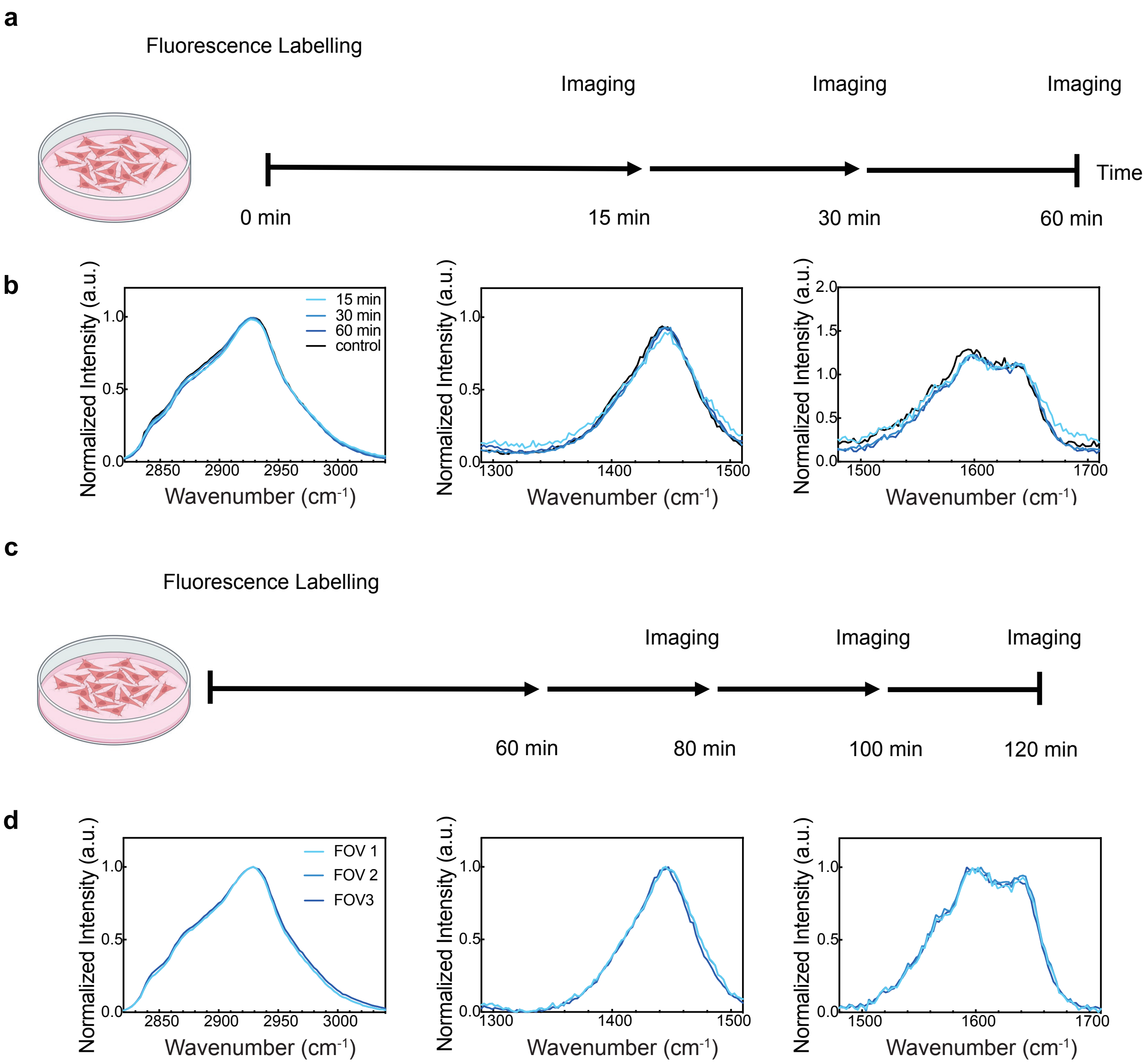


**Extended Data Figure 4: MitoTracker staining across variable durations do not perturb mitochondrial spectral signatures.**

(a) Experimental design used to assess potential effects of MitoTracker labeling time on mitochondrial measurements. (b) Averaged mitochondrial spectra acquired after different labeling durations, shown in the C-H window (left) and fingerprint window (1300-1500 $cm^{-1}$, center; 1500-1700 $cm^{-1}$, right). (c) Schematic of the time-lapse imaging protocol used to evaluate potential time-dependent perturbations during acquisition. (d) Mean mitochondrial spectra measured after varying time-lapse imaging durations, shown in the C-H window (left) and fingerprint window (1300-1500 $cm^{-1}$, center; 1500-1700 $cm^{-1}$, right).

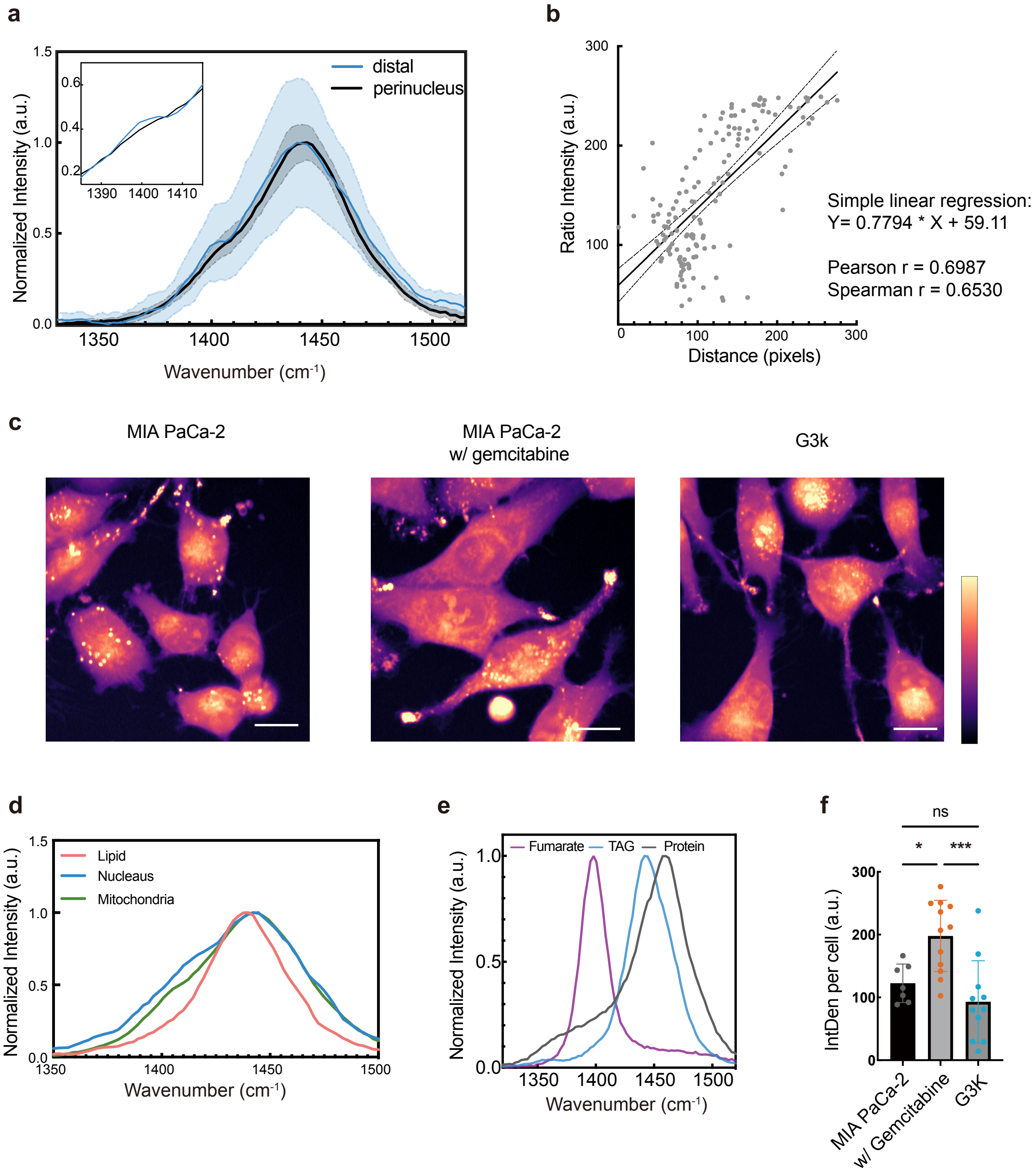

**Extended Data Figure 5: Subcellular spectral analysis reveals drug-induced fumarate accumulation in cancer cells.**

(a) Normalized SRS spectra of mitochondrial subgroups corresponding to Fig. 3b. Solid lines and shaded regions indicate mean ± SD. (b) Correlation between ratio-metric mitochondrial intensity and distance to the nuclear mass center in cisplatin-treated T24 cells (n = 149 mitochondria). Solid line, linear regression fit; dashed lines, 95% confidence interval. Pearson and Spearman correlation analyses were both performed. (c) Representative mean SRS image in the C-H window (2800-3000 $cm^{-1}$) for parental MIA PaCa-2 cells, gemcitabine-treated cells, and gemcitabine-resistant G3K cells (as shown in Fig. 3f). Scale bar: 15 μm. (d) Average hSRS spectra from ROIs corresponding to lipid droplets (n = 16), nucleus (n = 12), and mitochondria (n =

28). (e) Normalized reference spectra used for LASSO spectral unmixing. Fumarate, lipid, and protein components were represented by a 500 mM fumarate aqueous standard, triglyceride (-$CH_2$) and bovine serum albumin (-$CH_3$) respectively. (f) Per-cell fumarate signal quantified by integrated density for MIA PaCa-2 cells without gemcitabine (n = 7), with gemcitabine (n = 12), and G3K cells (n = 11) (one-way ANOVA with Tukey's multiple-comparison test; bars show mean ± SD; *P < 0.05, ***P < 0.001).

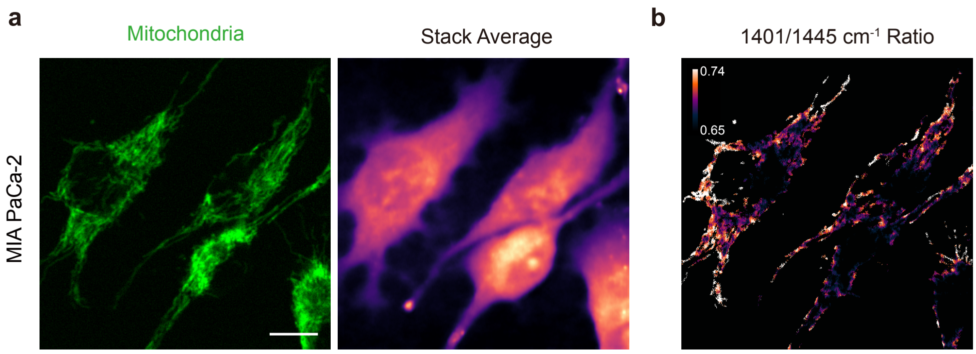


**Extended Data Figure 6: Ratio-metric analysis for mitochondria metabolism in MIA PaCa-2 cells under gemcitabine treatment.**

(a) Two-photon fluorescence image of MitoTracker-labeled mitochondria (left) and the corresponding hSRS stack average image (right). Scale bar: 15 μm. (b) Color coded ratio-metric mapping for mitochondria area with the same FOV in a.

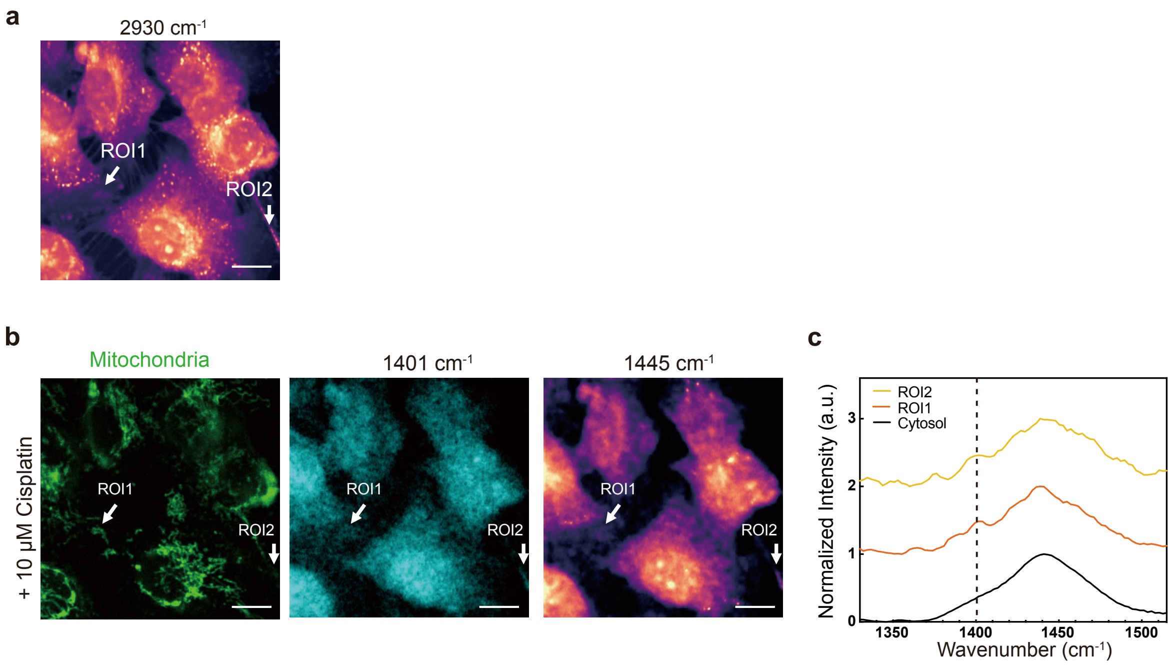


**Extended Data Figure 7: Spatially localized fumarate accumulation in microtube-associated mitochondria following chemotherapy.**

(a) Representative C-H window SRS image for T24 cells after cisplatin treatment. (b) Two-photon fluorescence image of MitoTracker-labeled mitochondria (left) and corresponding SRS images at 1401 $cm^{-1}$ (fumarate, middle) and 1445 $cm^{-1}$ ($CH_2$ deformation, right). Scale bar: 15 μm. (c) Normalized SRS spectra from ROI regions indicated by white arrows in b. Spectra are vertically offset for clarity.

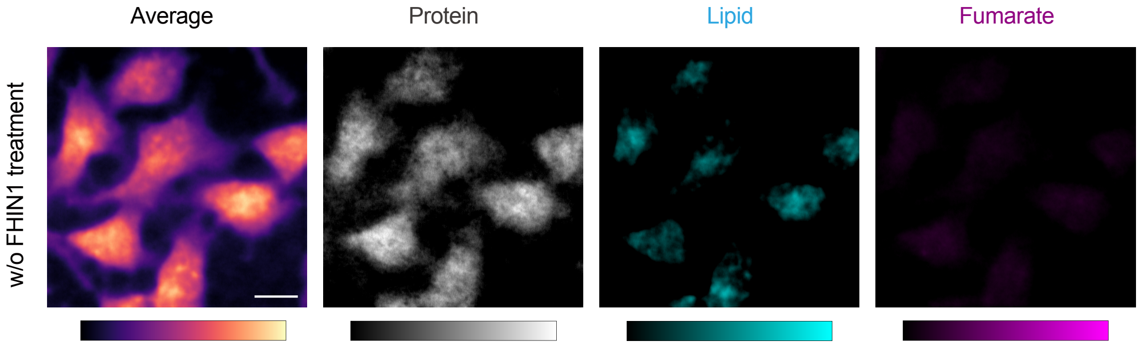


**Extended Data Figure 8: LASSO spectra unmixing validation.**
Representative hSRS average, LASSO unmixing results for hSRS images (channels are protein, lipid, and fumarate in sequence) for cells without FHIN1 treatment. Channels are in the same contrast with Fig 3.f.

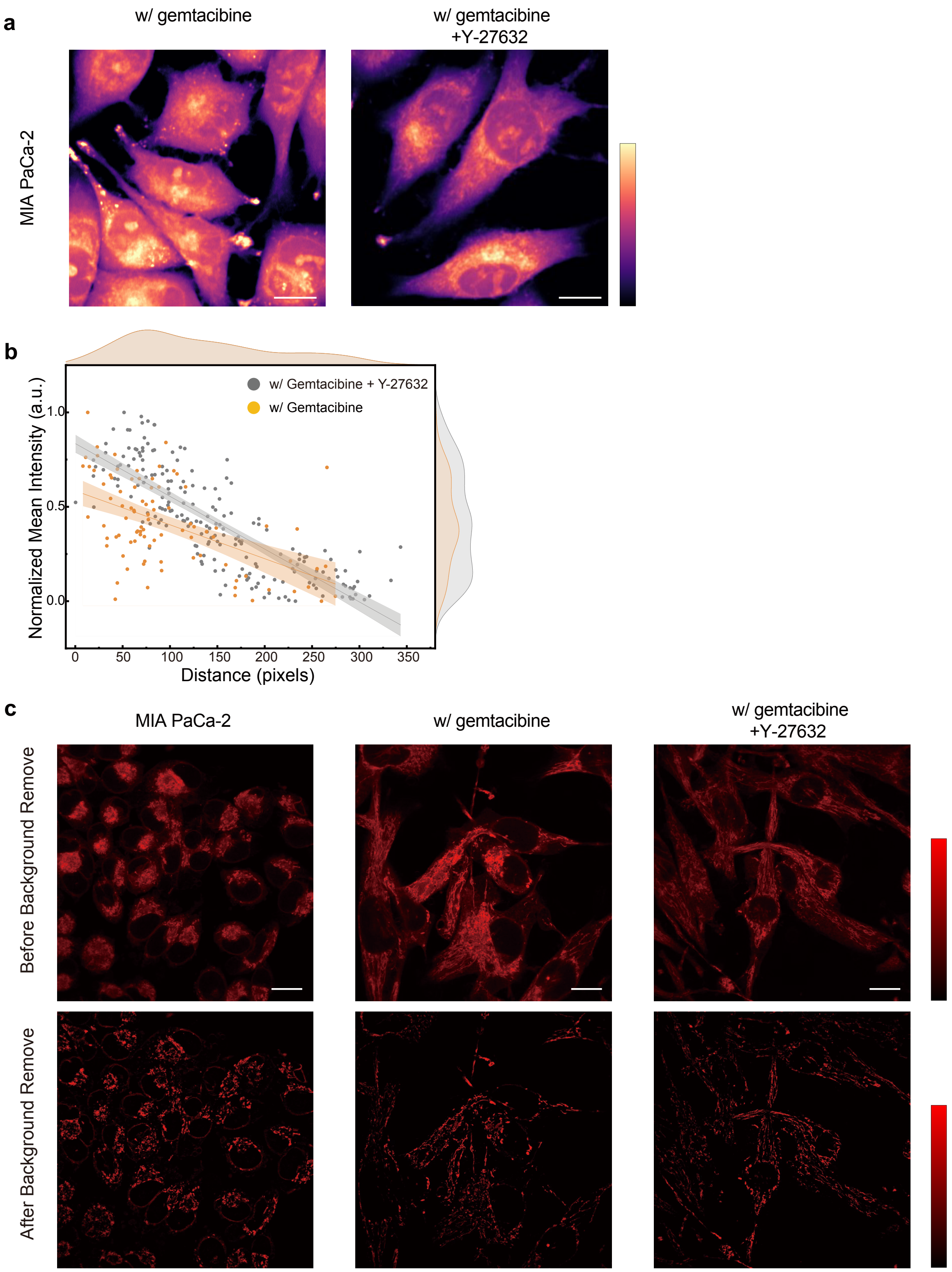


**Extended Data Figure 9: Mitochondria morphology changes under pharmacological stress.**

(a) Representative mean SRS image in the C-H window (2800-3000 $cm^{-1}$) for MIA PaCa-2 with or without Gemcitabine and G3k showed in Fig4 a. Scale bar: 15 μm. (b) Correlation between normalized mean fumarate intensity per mitochondrion and distance to the nuclear mass center in gemcitabine-treated MIA PaCa-2 cells without (n = 91) or with (n = 197) ROCK inhibitor. Solid line, linear regression; shaded band, 95% confidence interval. (c) Confocal fluorescence images of mitochondria labeled with MitoBrilliant 646 shown before and after background subtraction. Scale bar, 20 µm.

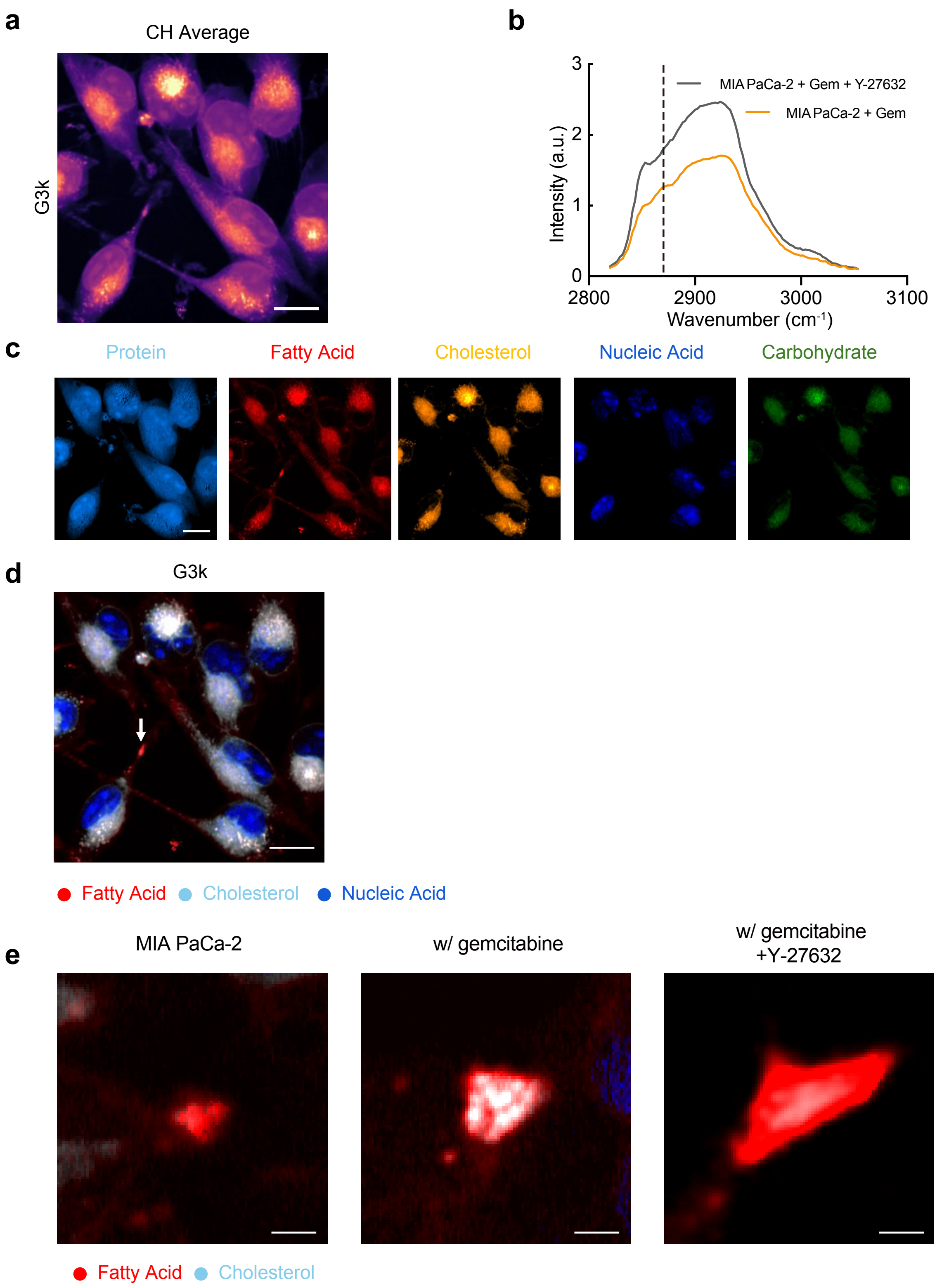


**Extended Data Figure 10: Lipid Droplets accumulation in the protrusions under stress.**

(a) Representative CH window average for G3k cells. Scale bar: 15 μm. (b) Raw SRS spectra acquired from LDs located in protrusions in the C-H region for gemcitabine-treated MIA PaCa-2 cells with or without Y-27632. (c) LASSO unmixing results for G3k cells (channels are fatty acid, cholesterol, nucleic acid and carbohydrate in sequence.

Scale bar: 15 μm. (d) Merged image of TAG, cholesterol and Nucleic Acid from LASSO results for G3k cells. Scale bar: 15 μm. (e) Zoomed in images in the dashed box in the fig. 5e. Scale bar: 2.5 μm.
